# Exploring fern pathosystems and immune receptors to bridge gaps in plant immunity

**DOI:** 10.1101/2025.03.18.643875

**Authors:** Baptiste Castel, Madeleine Baker, Jean Keller, Yves Martinez, Maxime Bonhomme, Pierre-Marc Delaux, Christophe Jacquet

**Affiliations:** Laboratoire de Recherches en Sciences Végétales, CNRS, Université Toulouse III Paul Sabatier, Institut National Polytechnique Toulouse, Castanet-Tolosan 31320, France; Fédération de Recherche 3450, Plateforme Imagerie, Pôle de Biotechnologie Végétale, Castanet-Tolosan 31320, France

**Keywords:** evoMPMI, fern, NLR, filamentous pathogens, gametophyte/sporophyte

## Abstract

Land plants include angiosperms, bryophytes, lycophytes and ferns, each of which may deploy distinct strategies to resist pathogens. Here, we investigate fern-pathogen interactions by characterizing novel pathosystems and analysing the diversity of fern immune receptors.

A collection of fern species was inoculated with a diverse set of microbes, and disease symptoms were assessed. We further leveraged published genome mining tools to analyse the diversity of RECEPTOR-LIKE KINASES, RECEPTOR-LIKE PROTEINS (RLKs/RLPs) and NUCLEOTIDE-BINDING AND LEUCINE-RICH REPEATS (NLRs), along with key immune signalling components, in ferns.

Our results reveal that ferns exhibit a range of responses to pathogens, including non-host resistance and specific resistance mechanisms. Among ten ferns tested, *Pteris vittata* displays the broadest spectrum of pathogen compatibility. Genome mining indicates that ferns encode a diverse repertoire of putative immune receptors, antimicrobial peptides and mediators of systemic acquired resistance. Ferns possess numerous RLKs/RLPs, resembling those required for cell-surface immunity in angiosperms. They also encode diverse NLRs, including a previously undescribed class with a disordered N-terminal domain, termed disN-NLR.

These findings provide insights into disease resistance evolution and open promising perspectives for crop protection strategies.

## INTRODUCTION

Ferns is an ancient plant lineage, with some fossils estimated to be 400 million years old (Devonian) (Nitta *et al*., 2022; Ali *et al*., 2024). But in fact, Polypodiales, which include about 80% of extant fern species, co-diversified with angiosperms around 100 million years ago, during the Cretaceous (Schneider *et al*., 2004). This suggests that ferns have experienced selective pressures similar to those of flowering plants. Yet their genomic evolution and environmental adaptations remain largely understudied. While angiosperm genomes have been extensively characterised, only a few fern genomes are publicly available. This includes the aquatic ferns *Azolla filiculoides* (Li *et al*., 2018), *Salvinia cucullata* (Li *et al*., 2018), *Marsilea vestita* (Rahmatpour *et al*., 2023) and *Ceratopteris richardii* (Marchant *et al*., 2022), the terrestrial ferns *Adiantum capillus-veneris* (Fang *et al*., 2022) and *Adiantum nelumboides* (Zhong *et al*., 2022), and the tree-fern *Alsophila spinulosa* (Huang *et al*., 2022). Fern genomes stand out for their large size, high chromosome numbers, long introns and abundance of repetitive sequences.

Historically, ferns have been primarily studied through morphological traits (Tryon, 1952; Christenhusz & Chase, 2014). While Linnaeus originally described 11 fern genera, recent phylogenies classify the ca 10,000 known species into 200-350 genera and 11 orders (Christenhusz & Chase, 2014; Nitta *et al*., 2022). Among these, Polypodiales represent 80% of all fern species, Cyatheales includes most tree ferns, Salviniales are aquatic and Equisetales form the sister lineage to all other ferns. Ferns form a monophyletic group with spermatophytes (seed plants) with which they share vascular tissues and branched leaf veins, yet remaining distinct due to their macroscopic gametophytic phase and lack of seed reproduction (Donoghue *et al*., 2021).

Ferns have been assumed to be less affected by pathogens than angiosperms, but this perception may reflect a bias towards angiosperm research (Antonovics, 2020). In the United States of America alone, at least 746 filamentous pathogens have been reported on ferns, including ascomycetes such as *Colletotrichum* (Jones *et al*., 2003) and *Fusarium* species (Mali *et al*., 2016), basidiomycetes such as *Puccinia* and *Hyalopsora* species, and oomycetes such as *Pythium* and *Achlya* species (Antonovics, 2023). These genera contain major pathogens of angiosperm crops (Cannon *et al*., 2012; Okungbowa & Shittu, 2012), suggesting that ferns could serve as a valuable genetic reservoir for disease resistance in crops.

Understanding how ferns defend themselves against pathogens could reveal both conserved and novel plant immunity mechanisms. In angiosperms, pathogen surveillance is mediated by both cell-surface and intracellular immune receptors (Jones & Dangl, 2006). Cell-surface immune receptors are generally receptor-like kinase (RLK) or receptor-like proteins (RLP), recognize conserved pathogen-associated molecular patterns (PAMPs) and activate PAMP-triggered immunity (PTI) (Bender & Zipfel, 2023). Intracellular immune receptors are generally nucleotide-binding and leucine-rich repeat (NLR), detect pathogen effectors and activate effector-triggered immunity (ETI) (Contreras *et al*., 2023). Three major classes of NLRs are: toll, interleukin1 and some *R*-genes product (TIR-NLRs), coiled-coil (CC-NLRs) and resistance to powdery mildew8 (RPW8-NLR). Bryophytes additionally contain lineage-specific kinase (Kin-NLRs and αβ-hydrolase (Hyd-NLRs) (Shao *et al*., 2016; Liu *et al*., 2023; Castel *et al*., 2024). PTI and ETI cooperatively enhance plant resistance to pathogens (Ngou *et al*., 2022a; Jones *et al*., 2024).

In this study, we investigated fern pathosystems by testing cross-compatibility between various ferns and microbes, including major crop pathogens. We also explored the diversity of immune receptors in ferns to assess their potential as a genetic reservoir for disease resistance. Our results showed that some crop pathogens, such as *Sclerotinia sclerotiorum* and *Fusarium proliferatum*, can infect multiple fern species. Among them, the fern *Pteris vittata* appears to be a particularly suitable model for studying fern immunity, as it displays compatibility with a broad range of tested microbes. Surprisingly, we found that gametophytes of *Pteris vittata* exhibit distinct resistance levels to certain pathogens compared to sporophytes. Additionally, we identified a diverse repertoire of immune receptors in ferns, including well characterised TIR-NLRs, CC-NLRs and RPW8-NLRs, but not the bryophyte-specific Kin-NLRs and Hyd-NLRs. Furthermore, we detected also non-canonical NLRs and described a previously unrecognized NLR subclass with a disordered N-terminal domain, which we termed disN-NLR. These results highlight the potential of ferns as a resource to explore plant immunity and improve disease resistance in crops.

## MATERIALS AND METHODS

### Fern and microbe material used in this study

A collection of fern species and microbial isolates (**Table S1**) was assembled for a systematic evaluation of cross-compatibility. Some fern specimens were obtained from external laboratories, others were collected from the wild, and some were purchased from commercial sources. Most ferns used here belong the *Polypodiales* order, the largest of ferns. Additionally, three species from the *Equisetales* order, the sister lineage to all other ferns were included (**Figure S1**). Only *Pteris vittata* could be successfully maintained and propagated under our laboratory conditions, while commercial ferns did not survive long-term. However, we still have access to the ferns harvested from the wild, due to their known geographic sampling area (Castanet-Tolosan, FR).

For microbial isolates, some were isolated from the bryophyte *Marchantia polymorpha* (Nelson *et al*., 2018) while others belong to our institutional collection (**Table S1** and **Figure S1**). These include broad-spectrum plant pathogens such as the ascomycete fungi *Sclerotinia sclerotiorum* (Purdy, 1979), *Colletotrichum spp.* (many of them were kindly provided by Dr Richard O’Connell, BIOGER, INRAE, Versailles-Saclay, France) (Cannon *et al*., 2012) or *Fusarium spp.* (Okungbowa & Shittu, 2012; Redkar *et al*., 2021), as well as host-specific pathogens like the oomycete *Aphanomyces euteiches* (specialist of Fabaceae) (Gaulin *et al*., 2007), and fungi not commonly considered pathogenic, such as *Biscogniauxia mediterranea*, *Coniochaeta sp.* and *Hypoxylon sp.* (Nelson *et al*., 2018).

### Plant growth conditions

Ferns at the sporophytic stage were grown in pots containing compost and pozzolan (3:1) in a greenhouse supplemented with LED, maintaining a 16-hour light / 8-hour dark photoperiod. Temperature was set at 22 °C and relative humidity at 70 %. Gametophytes were grown from surface-sterilized spores. Spores were sterilized as followed: 2 hours imbibition in water, 10 minutes in 1% bleach and 0.01% Tween-20, followed by three washes with sterile water. Spores were then sown on LS ½ medium [2.36 g/l (Caisson, LSP03), 0.8 g/l agar] in Petri dishes. Cultures were sealed with parafilm and maintained in a Snijder growth cabinet (type EB1) under 15-hour light / 9-hour dark photoperiod at 25 °C.

### Microbe culture and inoculation

Microbes were propagated on Potato Dextrose Agar (PDA, 39 g/l) at 22 °C in the dark. For long-term storage, spores were kept in 30 % glycerol at-80 °C. For mid-term storage, mycelium was stored in the dark at 10 °C on PDA Petri dishes. For inoculation, plugs of ca 5mm x 5mm containing actively growing mycelium were excised from 10-day-old cultures. For sporophytes, detached leaves (or stems for *Equisetum* species) were placed on a moist tissue towel. Gametophytes were inoculated directly on their growth media, approximatively two months after sowing, prior to sporophyte formation. PDA plugs with mycelium were placed in direct contact with the leaf or gametophyte surface, and the Petri dishes were sealed with parafilm to maintain high humidity. All collected ferns were inoculated in several trials with multiple pathogens. Disease symptoms were assessed at 8 days post inoculation (dpi) by measuring the area of brown tissue surrounding the inoculum site, using ImageJ software.

### Microscopy

Detached leaves or gametophytes were inoculated with *F. proliferatum*, *F. oxysporum* or *C. nymphaeae*, using 10 µl of a solution adjusted to 10^5^ spores/ml. Samples were collected at different infections stages for microscopic observations: spore germination (2 dpi) and mycelium colonization (6-17 dpi). For confocal microscopy, inoculated leaves were harvested and embedded in 7 % agarose in 6-well plates. Cross sections (100 µm thick) were obtained using a Leica VT1000S vibratome and incubated for 15 minutes in 1X Phosphate Buffer Saline (PBS) with 0.01 % calcofluor and 1 mg/l of Wheat Germ Agglutinin-AlexaFluor-488 (WGA-AF488). Cross sections were imaged using a Leica SP8 confocal microscope, with excitation at 488 nm and emission settings of 415-456 nm (calcofluor), 497-547 nm (WGA-AF488), and 700-750 nm (chloroplast autofluorescence). For scanning electron microscopy (SEM), 1×1 cm sections of leaves were fixed in a solution of 0.05 M sodium cacodylate buffer (pH 7.2) containing 2.5 % glutaraldehyde, dehydrated through an ethanol series, and critical-point dried using liquid CO₂. Samples were mounted on the observation plate using conductive silver paint and sputter-coated with platinum. Images were acquired using a Quanta 250 FEG FEI scanning electron microscope at 5 kV, with a spot size of 3 and a working distance of 10 mm.

### NLR and RLK/RLP mining

Immune receptors were predicted across available fern genomes and representative species from other land plant lineages (**Table S1**). To identify putative cell-surface immune receptors, we extracted RLKs and RLPs using RGAugury v2.2 (Li *et al*., 2016) which were further classified based on domain composition (**Table S2**). For intracellular immune receptors, NLRs were identified using both RGAugury and NLRtracker v0fe62b3 (Kourelis *et al*., 2021), followed by functional domain annotation with HMMER v3.2.1 to confirm the presence of NB-ARC domains. A subset of 3,775 NLRs (from 3,637 proteins) was retained for phylogenetic analysis. Protein sequences were aligned using MUSCLE v5.1.0 and trimmed with trimAl v1.4.1 to remove poorly aligned positions. Maximum-likelihood phylogenies were reconstructed using IQ-TREE v2.2.2.6, with the best-fitting evolutionary model selected via ModelFinder and branch support assessed with 10,000 replicates of both sh-aLRT and ultrafast bootstrap. The resulting phylogeny was visualized and annotated with iTOL v7. NLRs were classified into functional groups based on domain composition, and novel clusters were identified, including a previously undescribed class of NLRs with a disordered N-terminal domain, named disN-NLRs. Additional methodological details are provided (**Method S1, Data S1, S2 and S3**).

## RESULTS

### Ferns are hosts for various pathogens

We assessed the compatibility between diverse ferns and microbes (**Table S1** and **Figure S1**). Disease symptoms were quantified as the area of brown tissue, eight days after inoculation (**Figure 1**). Several microbes did not induce visible symptoms in any of the ferns tested. These include microbes that are not pathogenic on their natural host: *Hypoxylon sp.*, *Biscogniauxia Mediterranea*, *Coniochatea sp.* (Nelson *et al*., 2018). They also include host-specific pathogens, such as *Aphanomyces euteiches* and *Colletotrichum magnum*, suggesting non-adapted isolates or non-host resistance in ferns (Panstruga & Moscou, 2020). *Colletotrichum trifolii* race FR was also not compatible with any fern tested, but race 2 was able to grow on *Polystichum setiferum* and *Pteris vittata* (**Figure 1B**). This suggests that the lack of symptoms observed for race FR is more likely due to a specific resistance mechanism in these species, rather than non-host resistance. In contrast, *Sclerotinia sclerotiorum* and *Fusarium proliferatum* triggered significant disease symptoms in multiple fern species and exhibited the broadest host range among the pathogens tested. Among the ferns, *Pteris vittata* was susceptible to nearly all tested microbes, whereas *Nephrolepis exaltata*, *Polystichum setiferum*, and the three *Equisetum* species tested demonstrated broad resistance. Overall, these findings indicate that some generalist pathogens, including those known to infect angiosperms, can be compatible with ferns. Among the ferns, *P. vittata* appears to be broadly compatible with many microbes.

**Figure 1:**
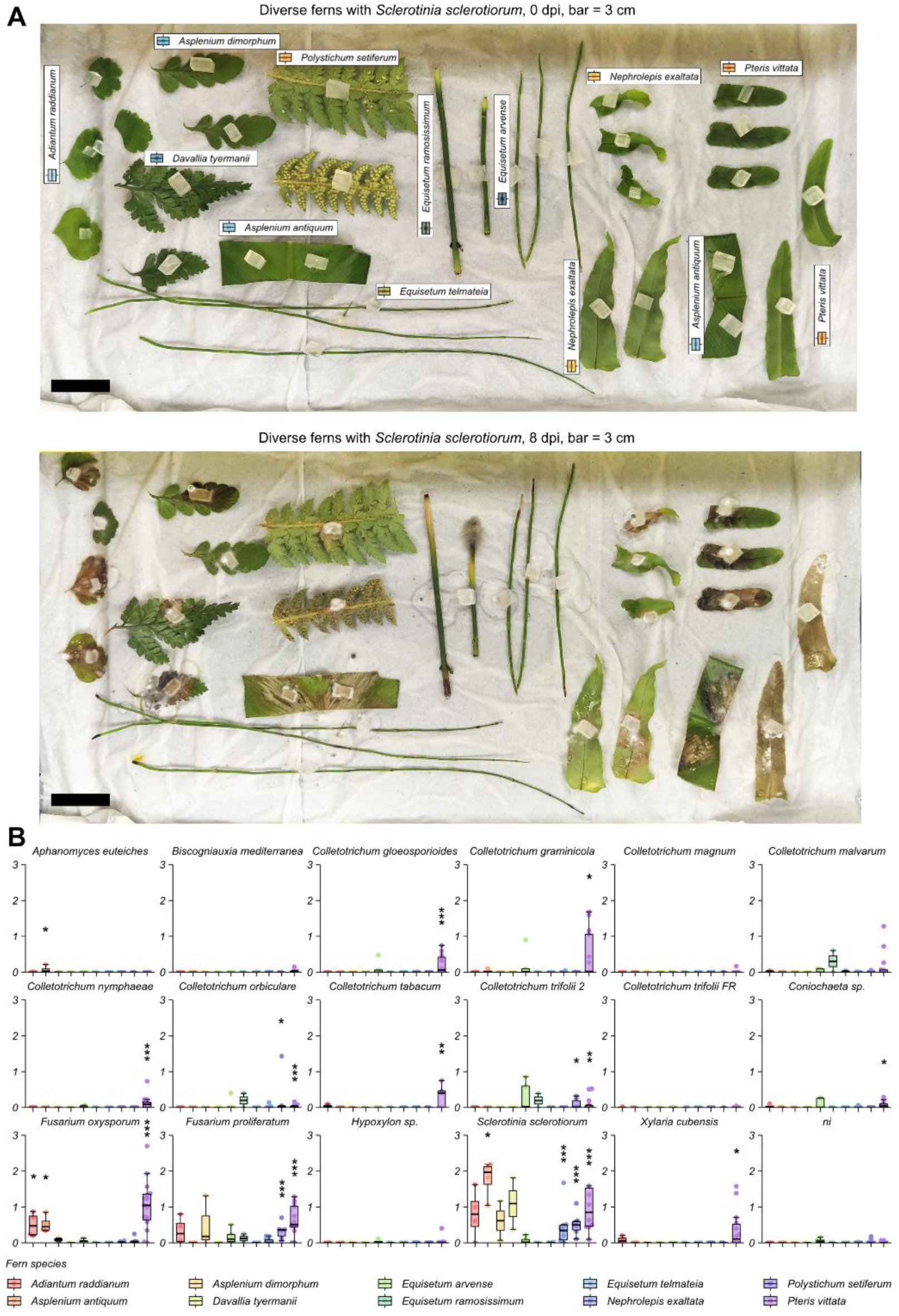
Some microbes cause disease symptoms in ferns (sporophyte) **A.** Representative disease symptoms caused by *Sclerotinia sclerotiorum* on diverse ferns. Detached leaves were inoculated with plugs of PDA medium containing actively growing fungus. Photos were taken at 0 dpi (top panel) and 8 dpi (bottom panel). Fern species names are indicated in the top panel. Bar = 3 cm **B.** Quantification of disease symptoms caused by microbes on ferns. Leaves (or stem for *Equisetum spp.*) were collected and placed on a moist paper towel as in panel ‘A’. Tissues were inoculated with PDA plugs media containing actively growing microbes. Disease symptoms were measured at 8 dpi, as the area (in cm²) of brown tissue extending from the inoculation sit, using ImageJ. Due to limited plant material, the number of replicates (n) ranges from 2 to 15 (average ∼5) across independent inoculations. *, ** or *** indicate p-value <0.05, <0.01 or <0.001 respectively, from a t-test for normally distributed data or a Wilcoxon test for non-normally distributed data (normality was assessed using a Shapiro test), for values with a mean >0.05 cm², as smaller values are too smalls to reliably describe symptoms. Figure generated with the R package ‘tidyplots’.

### Gametophytes and sporophytes respond differently to some pathogens

The fern life cycle consists of two morphologically and functionally distinct phases: the haploid gametophyte, which produces gametes via mitosis, and the diploid sporophyte, which develops after fertilization and represents the dominant life stage in most species (Watkins *et al*., 2007a). Because sporophytes and gametophytes differ in structure and physiology, particularly in the presence of vascular tissues, cuticle development, and overall size, we tested whether they could respond differently to the same pathogens (**Figure 2**).

**Figure 2:**
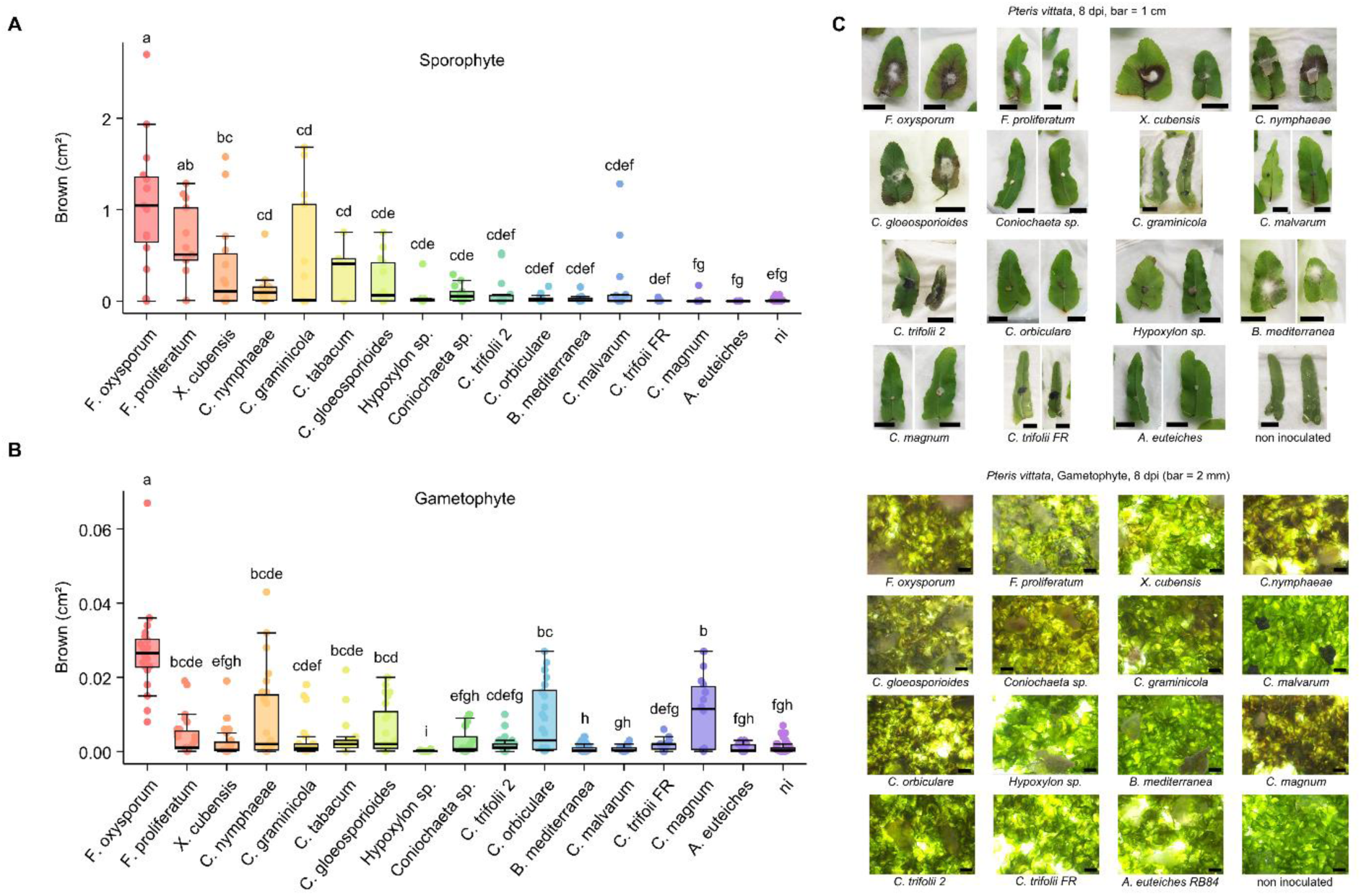
Sporophytes and gametophytes of *Pteris vittata* respond differently to some pathogens A.B. Quantification of disease symptoms on sporophytes (**A**) or gametophytes (**B**) of *Pteris vittata*. Disease symptoms were measured as the area (cm²) of brown tissue extending from the inoculation site at 8 dpi. Letters indicate significantly different groups, analysed independently for each plot A and B, based on an ANOVA followed by a Duncan test on data normalized by quantiles, with significant threshold p-value <0.05. Plots were generated with the R package ‘tidyplots. **A.** Detached leaves of sporophytes were inoculated with plugs containing a given microbe. Due to limited plant material, the number of replicates (n) ranges from 5 to 15 (average ∼13), across independent inoculations. **B.** Gametophytes of *Pteris vittata* were grown under sterile conditions and inoculated with plugs containing a given microbe. Due to limited plant material, the number of replicates (n) ranges from 17 to 36 (average ∼27), across independent inoculations. **C.D.** Representative images of symptoms in *Pteris vittata* sporophytes (**C**, bar = 1 cm) and gametophyte (**D**, bar = 2 mm). Photos were taken at 8 dpi.

Pathogen inoculations revealed distinct susceptibility patterns in *P. vittata* sporophytes (**Figure 2A** and **C**). Two tested *Fusarium* species were highly aggressive, inducing strong symptoms that initially localized in the veins before spreading to surrounding leaf tissues. Similarly, *Xylaria cubensis*, a natural pathogen of *Marchantia polymorpha* (Nelson *et al*., 2018), generated large necrotic and dead tissue areas, suggesting an active necrotrophic behaviour. A second group of pathogens, including *C. nymphaeae*, *C. graminicola* and C*. tabacum*, triggered maceration symptoms but in a less frequent or less extensive manner. Finally, the remaining microbes exhibited rare, weak or no visible pathogenic effects, failing to induce significant symptoms across different assays.

Gametophytes exhibited lower symptom levels compared to sporophytes, suggesting a slower progression of fungal colonization (**Figure 2B** and **D**). Moreover, the relative susceptibility to different microbes did not follow the same ranking as in sporophytes. *F. oxysporum, F. proliferatum*, *C. nymphaeae* and *C. tabacum*, which had already exhibited strong pathogenicity in sporophytes, also caused significant symptoms in gametophytes. In contrast, some fungi that successfully colonized sporophytes failed to invade gametophyte tissues. This was particularly evident for *C. graminicola* and *X. cubensis*, which caused severe symptoms in sporophytes but were unable to establish infection in gametophytes, that generally remain green. Conversely, some pathogens, that failed to induce symptoms in sporophytes caused significant browning in gametophytes, such as *C. orbiculare* and *C. magnum*.

These findings suggest that the pathogen virulence varies between gametophytes or sporophytes in *P. vittata,* highlighting a generation-dependent component of susceptibility.

### Pathogens can successfully colonize fern tissues

Among the different pathogens tested, *Colletotrichum nymphaeae*, *Fusarium proliferatum* and *Fusarium oxysporum* were selected for further microscopic analyses due to their virulence on both gametophytes and sporophytes in *Pteris vittata* (**Figure 2**). We investigated their infection processes in *P. vittata* using scanning electron microscopy (SEM), confocal laser scanning microscopy, and bright field/fluorescence microscopy (**Figure 3**).

**Figure 3:**
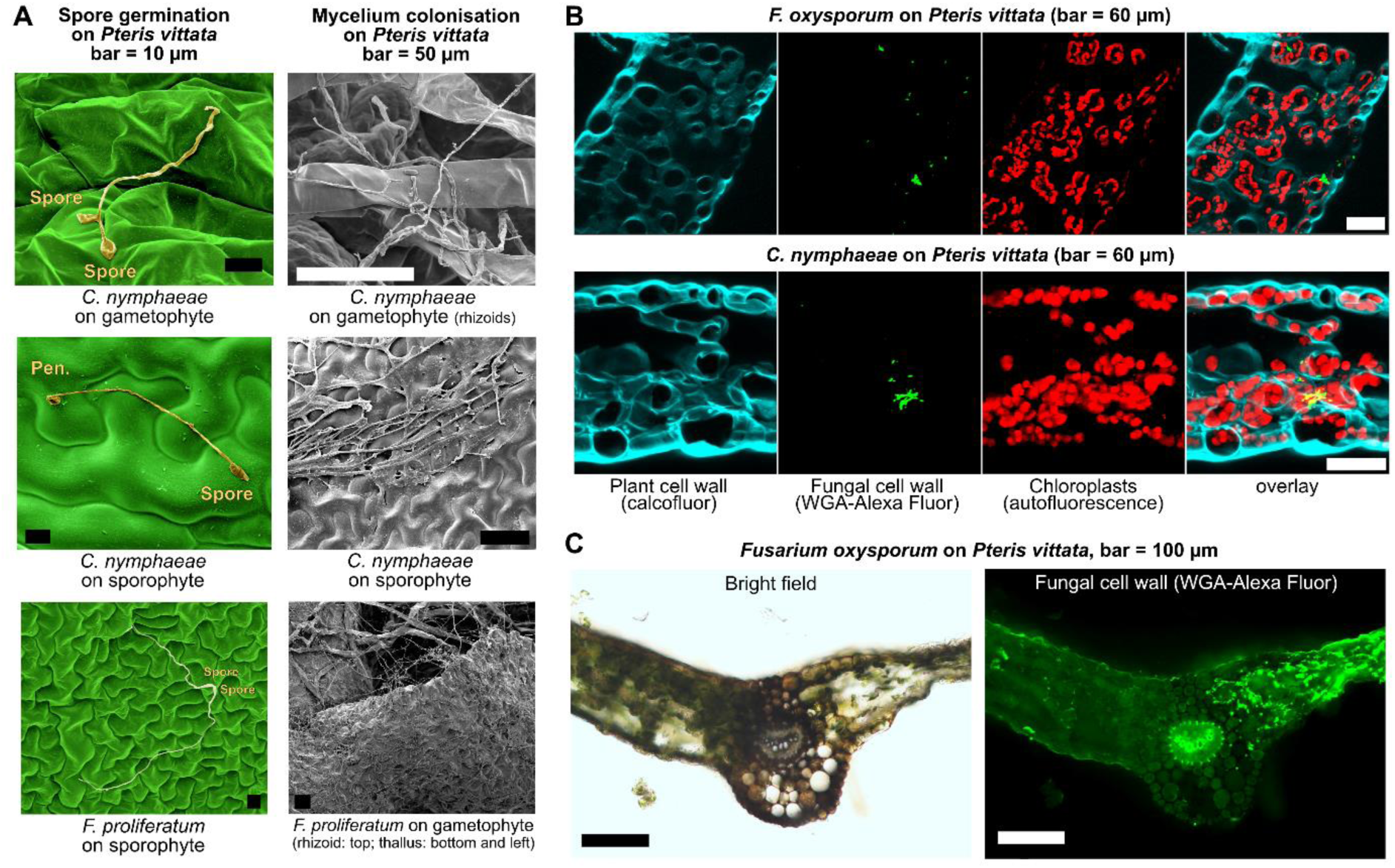
**Microbes can colonize superficial and internal tissues of *Pteris vittata* A.** Scanning electron microscopy (SEM) images showing fungal germination and development on the epidermis of *Pteris vittata* gametophytes and leaves. Left panels: False-coloured images (Affinity Designer v2.5.5), showing germinated spores and appressorium-like structures formed by *C. nymphaeae* at 2 dpi. Scale bar = 10 µm. Right panels: Later infection stages show extensive hyphal colonization of superficial tissues in *P. vittata* sporophytes or gametophytes at 6 dpi. Host cells in contact with mycelium appear swollen and macerated. Pen. = penetration structure (appressorium-like). Scale bar = 50 µm. **B.** Confocal microscopy images of 100 µm cross-sections leaves of *P. vittata* at the fungal migration front showing intercellular hyphal colonization with WGA-AlexaFluor (green). Plant cell walls are stained with calcofluor (blue). Autofluorescence of chloroplasts (red) indicates that plant cells are still alive at this stage. Picture taken at 17 and 6 dpi respectively. **C.** Microscopy images of 100 µm cross-sections leaves of *P. vittata* with *Fusarium oxysporum* showing extensive maceration and tissue disorganization on the right side of the leaf, with cell wall degradation, at four dpi. Left panel: bright field. Right panel: WGA-AlexaFluor staining (green) highlighs massive hyphal colonization in destructured tissues and intercellular fungal progression in the left-side tissue. Autofluorescence of lignin is visible in the central vasculature. Bar = 100 µm.

SEM images revealed that both *F. proliferatum* and *C. nymphaeae* germinated on the surface of *P. vittata* sporophyte and gametophyte tissues (**Figure 3A, left panels**). *C. nymphaeae* produced appressorium-like structures, suggesting a penetration strategy similar to that on waterlilies and strawberry (Johnson *et al*., 1997; Rose & Damm, 2024). No penetration structures were observed for *F. proliferatum*, suggesting a direct penetration through cell wall digestion. At later infection stage (6 dpi), hyphae formed dense networks across the surface of both life stages. Host cells in direct contact with the mycelium appeared swollen and macerated, consistent with tissue degradation observed in necrotrophic infections (**Figure 3A, right panels**).

To evaluate fungal penetration beyond surface colonization, we performed confocal microscopy on leaf cross-sections (**Figure 3B**). Sections performed in the fungal migration front of leaves inoculated with *C*.

*nymphaeae* (6 dpi) and *F. oxysporum* (17 dpi) exhibited intercellular hyphal growth, as visualized with WGA-AF staining, in living tissues, as indicated by the presence of numerous autofluorescent chloroplasts. No signs of intracellular hyphal colonization were observed at this stage, suggesting that both fungi primarily invade via intercellular spaces in *P. vittata*. Bright-field microscopy performed in symptomatic tissues confirmed severe tissue degradation in *P. vittata* leaves caused by *F. oxysporum* (**Figure 3C**), with extensive cell wall degradations and disorganized tissues, as well as massive colonization by fungal hyphae, visualized by WGA-AF fluorescence, likely indicating a switch to a necrotrophic fungal stage.

Together, these findings confirm that *P. vittata* is a viable host for *C. nymphaeae*, *F. proliferatum* and *F. oxysporum*, with successful pathogen attachment, penetration, and intercellular colonization. Many of the observed infection characteristics resemble those of plant filamentous pathogens showing a hemibiotrophic lifestyle in angiosperms (Perfect & Green, 2001).

### Fern genomes reveal a rich diversity of putative immune receptors

Receptor-like kinases (RLKs), receptor-like proteins (RLPs) and nucleotide-binding leucine-rich repeat receptors (NLRs) are major classes of immune receptors in angiosperms (Barragan & Weigel, 2020; Contreras *et al*., 2023). To determine whether similar receptors exist in ferns, we analysed available fern genomes, along with some control species representing the diversity of land plants (Table S1). We identified 11,390 RLKs (including 3,286 from ferns), 1,843 RLPs (including 371 from ferns) (Table S2) and 3,775 NLRs (including 396 from ferns) (Table S3).

### Fern genomes encode RLK/RLP

Within the RLK/RLP, we identified well-known receptor classes including leucine-rich repeat (LRR), lectin, malectin, LRR-Malectin (LRR-Mal.), wall-associated kinase (WAK), Thaumatin, cysteine-rich repeat (CRR), regulator of chromosome condensation 1 (RCC1) and lysin motif (LysM) (Table S2) (Lehti-Shiu *et al*., 2012; Dievart *et al*., 2020). Their distribution in ferns is comparable to that of other land plants, with LRR-and Lectin-RLKs being the most abundant among RLKs, and LRR-RLPs being the most abundant among RLPs (**Figure 4**). All categories are present in all fern genomes investigated.

**Figure 4:**
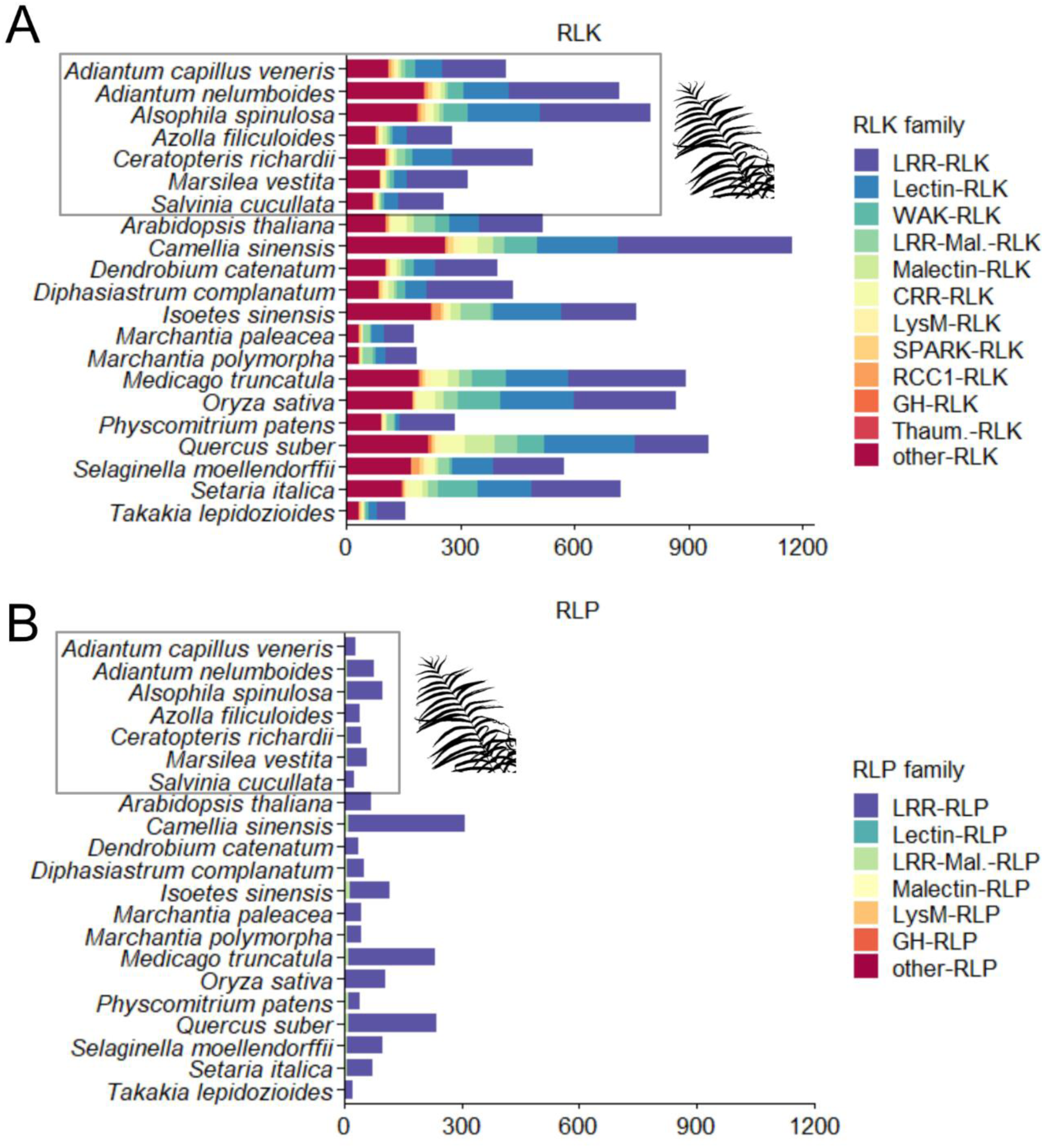
Diversity of cell-surface receptors in ferns. Distribution of receptor-like kinases (RLKs, **A**) and receptor-like proteins (RLPs, **B**) in ferns and other plant lineages. RLK and RLP were identified from plant genomes using RGAugury (10.1186/s12864-016-3197-x) and classified into receptor families based on HMM scan predictions. Plots were generated with the R package ‘tidyplots’.

In addition, some RLK/RLPs could not be assigned to any of these categories. We investigated whether they could represent fern-specific classes. The predicted domains were mostly identical between ferns and non-ferns unclassified RLK/RLPs (**Figure S2A**). We identified Romo1 (IPR018450), Vps39_1 (IPR019452), Photo_RC (PF00124) and DUF642 (PF04862) unique to ferns. However, we could identify some of these domains in tandem with kinases in angiosperms and/or gymnosperms such as A0A3P6FCZ3 (Vps39_1-kinase from *Brassica oleracea*), A0A438D9U7 (Photo_RC-kinase from *Vitis vinifera*) and A0AA38CBH3 (DUF642-kinase from *Taxus chinensis*), in the InterPro database (https://www.ebi.ac.uk/). This indicates that these classes of proteins are not specific to ferns.

### Fern genomes encode well-described NLRs

TIR-NLRs form three distinct clusters, with ‘Cluster I’ being the most widespread across land plants, including angiosperms, bryophytes, and ferns. ‘Cluster II’ is specific to ferns, while ‘Cluster III’ includes both ferns and bryophytes and is unexpectedly related to certain CC-NLRs from tea (*Camelia sinensis*), suggesting potential functional diversification. RPW8-NLRs, which are involved in immune signalling, are present across all land plant lineages. CC-NLRs segregate into three main groups: ‘EDVID’ and ‘G10,’ both specific to angiosperms, and ‘Cbl-N,’ which is found only in non-angiosperms, including ferns, lycophytes, and bryophytes. These findings suggest that CC-NLRs underwent lineage-specific expansions, with ‘EDVID’ and ‘G10’ evolving predominantly in spermatophytes, while ‘Cbl-N’ represents the ancestral state retained in non-angiosperms.

### Fern genomes encode a novel type of NLRs: disN-NLRs

We identified three additional clusters of fern NB-ARC-containing proteins: TIR-NB-TPRs, CC-NB-ARMs and one unclassified lineage (**Figure 5A**). TIR-NB-TPRs form a conserved class of NLRs, potentially regulating immunity, containing a TIR domain but signalling independently of TIR-NLRs (Johanndrees *et al*., 2023). CC-NB-ARMs are also conserved in plant genomes (Urbach & Ausubel, 2017; Kourelis *et al*., 2021). The rice CC-NB-ARM ‘RLS1’ could be a regulator of immunity (Wang *et al*., 2023).

**Figure 5:**
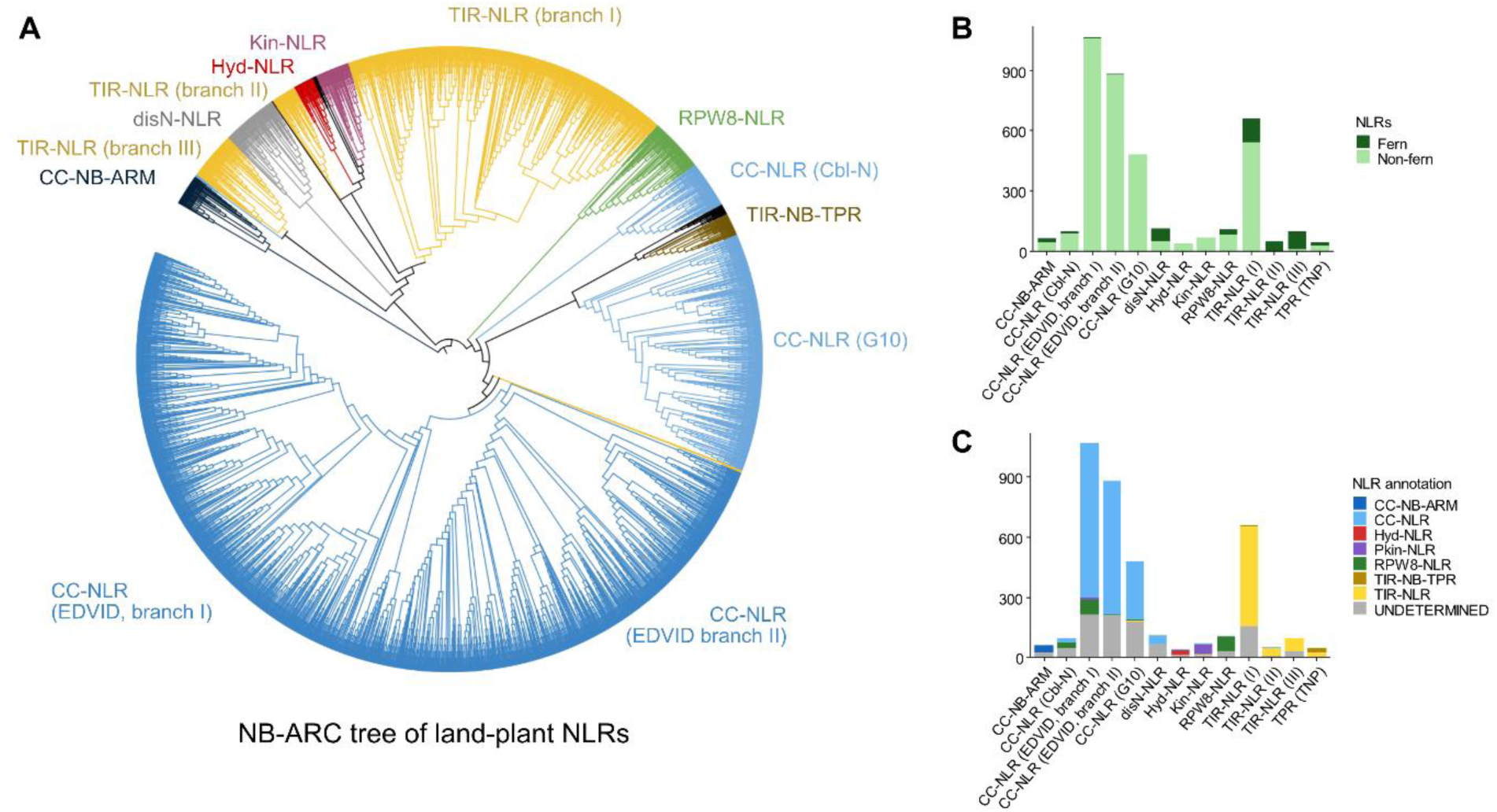
Diversity of NLRs in ferns and other land plants. **A.** Phylogeny of NB-ARC domain from NLRs across representative land plants, including ferns, mosses, liverworts, lycophytes, monocots and dicots (species detailed in Table S1). Some branches were manually coloured based on annotation. The indicated clades are supported by bootstrap values > 70/100. Branch lengths are not to scale. The tree on Newick format is available in Data S1 and its coloured information in Data S2. Tree visualization was generated with iTOL (https://itol.embl.de/). **B.** Distribution of fern NLRs across the different NLR clades. **C.** Repartition of NLR-type within each NLR clade, based on annotation from NLRtracker, RGAugury and/or HMM scan annotation. Plots in B and C were generated with the R package ‘tidyplots’.

The unidentified NLR branch is closely related to Hyd-, Kin-and TIR-NLRs clusters I and II (**Figure 5A**). It is annotated as OG5 and OG11 in previous studies (Chia *et al*., 2024). This cluster is particularly rich in fern NLRs (**Figure 5B**), prompting further investigation. Proteins from this cluster were sometimes classified as CC-NLRs, but mostly unclassified (**Figure 5C**). Two angiosperms NLRs are found in this cluster: MtrunA17Chr3g0097641 from *Medicago truncatula* and At4g09420 (or TN15) from Arabidopsis. MtrunA17Chr3g0097641 has two NB-ARC domains: one central clustering with ‘EDVID’ CC-NLRs, and one C-terminal clustering with disN-NLRs (**Data S1**). It suggests that it is a CC-NLR with a C-terminal NB-ARC integrated decoy (Cesari *et al*., 2014; Baggs *et al*., 2017; Marchal *et al*., 2022). TN15 is usually classified as a TIR-NB-ARC (Nandety *et al*., 2013), and could be accidentally clustered with disN-NLR in our analysis. Hence, this unclassified class of NLR could be specific to non-angiosperm land plants. We generated a wordcount plot of the HMM from this cluster (**Figure S2B**) but could not deduce a characteristic function. We used AlphaFold (Jumper *et al*., 2021) to predict the structures of representative full-length fern NLRs from this cluster (**Data S4** and **S5**), incorporating one molecule of ADP (**Figure 6A** and **Data S6**). ADP likely stabilizes the NB-ARC domain in its monomeric state (Tameling *et al*., 2006; Wang *et al*., 2019). AlphaFold modelized the ADP-bound NB-ARC and LRR domains with high confidence. However, the N-terminal domain consistently exhibited low structural confidence, often assembled as a four-helix bundle reminiscent of a CC domain, but with very low confidence and an extended disorder tail (**Figure 6**). It suggests that this N-terminal domains could be disordered, although it may adopt an ordered structure in some conditions. Therefore, we propose the name ‘disordered N-terminal NLR’, or disN-NLR for this novel class. The AlphaFold database (Varadi *et al*., 2024) contains a non-fern NLR from this cluster: Mp4g08790 from *Marchantia polymorpha*. Its predicted model shows well-structured NB-ARC and LRR regions, but a low confidence N-terminal domain, similar to fern disN-NLRs (**Figure 6B**).

**Figure 6:**
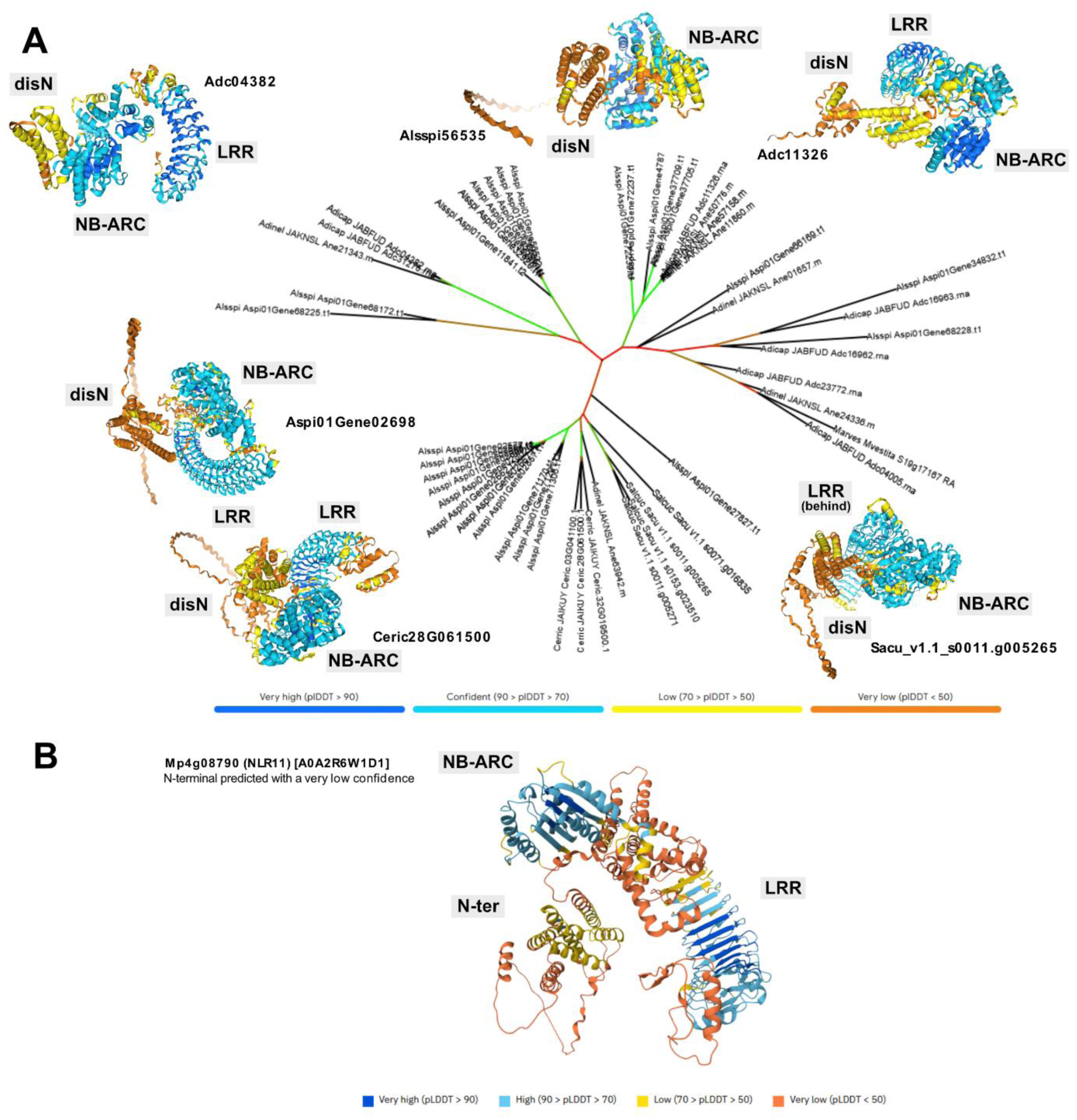
Fern disN-NLRs display an intrinsically disordered N-terminal domain. **A.** Phylogenetic tree of the N-terminal domain (aa 1-402) of fern disN-NLRs, with corresponding AlphaFold structural prediction of full-length proteins (Jumper *et al*., 2021). The colours of the branches indicate bootstrap values from 0 (red) to 100 (green). Protein structure confidence ranges from low (orange) to high (blue). The sequence alignment is available as Data S3 (FASTA format), the tree as Data S4 (Newick format) and the AlphaFold structures as Data S6. **B.** Predicted structures of a disN-NLR from *Marchantia polymorpha*, from the Alphafold database (Varadi *et al*., 2024). AlphaFold reference numbers are indicated in brackets next to the protein names.

## DISCUSSION

### Ferns are compatible with a broad range of pathogens

The literature on fern diseases is sparser than for angiosperms (Antonovics, 2020). Yet, several pathogens have been reported on ferns, primarily filamentous fungi (Stevenson, 1945; Toome *et al*., 2014; Kirschner *et al*., 2019; Antonovics, 2023), but also bacteria (Kloepper *et al*., 2013) and viruses (Sastry *et al*., 2019). Some of the reported filamentous pathogens belong to the same species as crop pathogens, such as *Colletotrichum gloeosporioides* (Jones *et al*., 2003) and *Botrytis cinerea* (Antonovics, 2023), both causing diseases on many crops including soybean, wheat and tomato (Sharma & Kulshrestha, 2015; Caseys *et al*., 2021).

Here, we expanded this category by identifying *Fusarium oxysporum*, *Fusarium proliferatum*, *Sclerotinia sclerotiorum*, *Colletotrichum nymphaeae* and a specific race of *Colletotrichum trifolii* as pathogens capable of colonizing ferns (**Figure 1**). This raised the question of whether these interactions reflect true compatibility or opportunistic infections facilitated by stress conditions in experimental settings. Microscopy observations revealed several conserved features of microbe pathogenicity (**Figure 3**). Pathogens such as *C. nymphaeae* and *F. oxysporum* retained classical infection strategies, including the formation of appressorium-like structures, intercellular hyphal development, and an alternation between biotrophic and necrotrophic phases. These observations align with previous findings on the evolutionary dynamics of plant-parasite interactions, which suggest that many core mechanisms of pathogen virulence remain conserved despite host diversification (Brown & Tellier, 2011).

The inability of *Aphanomyces euteiches* to infect any of the tested fern species tested suggests non-host resistance, a widely observed phenomenon in plant-pathogen interactions where an entire plant lineage remains incompatible with a given pathogen (Panstruga & Moscou, 2020). Conversely, the differential susceptibility observed between *P. vittata* and the two different tested races of *C. trifolii* strongly suggest host-race specific immunity, which may be linked to the recognition of an effector present in race FR, but not in race 2. The ferns pathosystems reported here provide valuable information to study the evolution of plant-pathogen interactions, offering new perspectives on resistance mechanisms in land plants.

### *Pteris vittata* could be a model to study fern-pathogen interactions

*Pteris vittata* exhibits the broadest range of compatibility with microbes, including pathogens that typically infect angiosperms or bryophytes (**Figure 1** and **2**). This fern is particularly interesting as it can accumulate arsenic (Ma *et al*., 2001; Gumaelius *et al*., 2004), likely through the arsenate reductase PvACR2 (Ellis *et al*., 2006), and it can establish arbuscular mycorrhiza symbiosis with the fungus *Glomus intraradices* (Martinez *et al*., 2012), making it a valuable model to investigate the conservation of the Common Symbiosis Signalling Pathway (CSP) in ferns (Radhakrishnan *et al*., 2020). In addition to its role in soil arsenic decontamination and symbiosis, we found that *Pteris vittata* is partially compatible with multiple pathogens, including *F. oxysporum*, *F. proliferatum*, *X. cubensis*, *C. nymphaeae* and *C. trifolii* race 2. These findings strengthen the potential of *P. vittata* as a model to study fern-microbe interactions and the evolution of molecular plant-microbe interactions.

### Gametophyte and sporophytes respond differently to pathogens

The life cycle of ferns consists of two independent and macroscopic phases: the gametophyte and the sporophyte (Christenhusz & Chase, 2014). Fern gametophytes, called prothalli, develop from spore germination. They are generally heart-shaped, single cell layer and few millimetres in size. They generate gametes, whose fertilization gives rise to a sporophyte, the dominant form of most ferns. The sporophyte forms roots, a stem (called rhizome, even when above ground) and leaves (called fronds) which bear spore-producing sporangia. Spore germination results in a new gametophyte. These structural and physiological differences suggest that sporophytes might be more resilient to stress. Surprisingly, our results revealed that *P. vittata* gametophytes can exhibit greater resistance to some fungi such as *Colletotrichum graminicola* and *Xylaria cubensis*, challenging the assumption that gametophyte are inherently more susceptible to pathogens (**Figure 2**).

Despite their seemingly fragile structure, gametophytes demonstrate remarkable resilience to environmental stresses. For instance, fern gametophytes exhibit a high tolerance to desiccation (Watkins *et al*., 2007b), a trait that is rare in sporophytes and likely a homoplasy (Proctor & Tuba, 2002; Farrar *et al*., 2008). Fern gametophytes also show adaptation to light stress (Krieg & Chambers, 2022), likely through the accumulation of reactive oxygen species scavenging molecules, such as β-carotene or tocopherols (Tausz *et al*., 2001; Fernández-Marín *et al*., 2012). Additionally, some fern gametophytes are also more resistant to freezing than their sporophytes, enabling their survival under snow during winter (Sato & Sakai, 1980). These differences suggest that gametophytes and sporophytes deploy distinct physiological or biochemical defence mechanisms. In angiosperms, developmental stage-dependent resistance to pathogens has been also reported, with mature plants generally more resistant than seedlings (Develey-Rivière & Galiana, 2007; DeMell *et al*., 2023). In the case of *P. vittata*, gametophytes appear more resistant than sporophytes to *C. graminicola* and *X. cubensis* (**Figure 2**). One possible explanation is the absence of vasculature in gametophytes, which may limit systemic pathogen spread. Another possibility is the accumulation of antimicrobial compounds specific to gametophyte. Indeed, ferns are known to produce a wide range of specialised metabolites in response to pests (San Francisco & Cooper-Driver, 1984; Page, 2002; Guha *et al*., 2005; Mitoi *et al*., 2024). Further investigation, particularly at the molecular level, is needed to determine whether gametophyte-specific immune strategies exist and how they compare to those of sporophytes

### Immune response may be conserved within tracheophytes

Tracheophytes have been exposed to pathogen infections for at least 407 million years (Strullu-Derrien *et al*., 2023), and likely since terrestrialization (Upson *et al*., 2018; Delaux & Schornack, 2021; Castel *et al*., 2024). Our findings indicate that ferns can resist to angiosperm pathogens, raising the question of whether these defence mechanisms are ancestral traits shared with seed plants or represent lineage-specific innovations. To investigate this question, we examined the conservation of well-characterized immune-related genes in ferns.

A recent study interrogated 26 fern genomes or transcriptomes (Ali *et al*., 2024), found that many immune genes are conserved in ferns (**Figure 7** and **Table S4**), including *BAK1* and *BIK1* required for signalling of many LRR-RKs (Bender & Zipfel, 2023). Two well-characterised CC-NLRs were not detected in ferns (**Figure 7**). However, our phylogenetic confirmed the presence in ferns of CC-NLRs, but only from the ‘Cbl-N’ type, which was lost in angiosperms (**Figure 5**). Despite their sequence divergence, ‘Cbl-N’ CC-NLRs likely functions similarly to ‘EDVID’ CC-NLRs found in angiosperms (Chia *et al*., 2024). In addition, RIN4, a protein targeted by multiple effectors and guarded by several CC-NLRs in angiosperms (Kim *et al*., 2022), is well conserved in ferns (**Figure 7**). This suggests that CC-NLR-mediated immunity is ancestral to land plants, but angiosperms evolved distinct ‘EDVID’ and ‘G10’ classes of CC-NLRs, while other lineages including ferns retain the ‘Cbl-N’ ancestral class.

**Figure 7:**
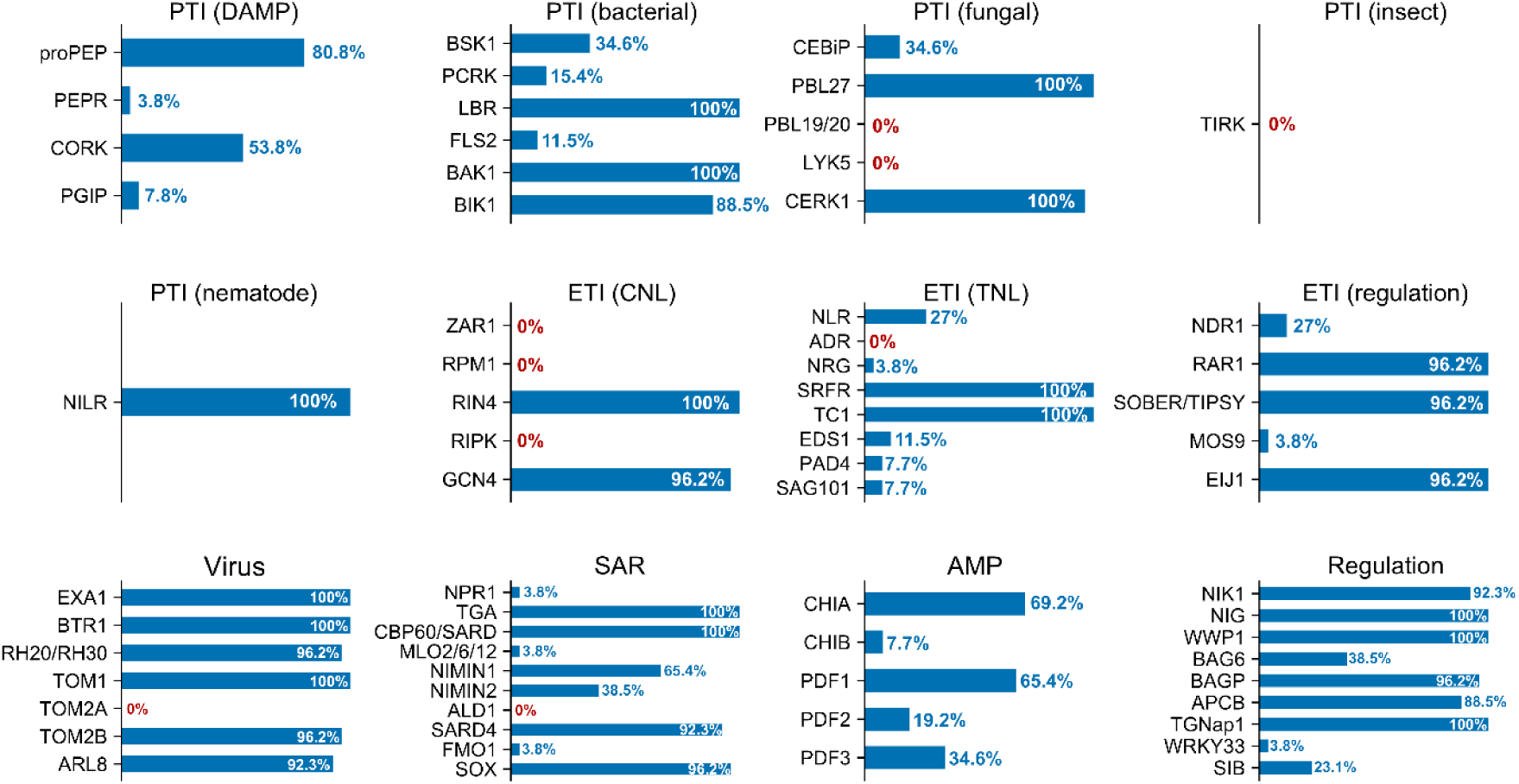
Prevalence of known immune-related genes across 26 ferns transcriptomes. Data extracted from (Ali et al., 2024, Table S9). Genes annotated as involved in response to pathogen or virus were included. The plot was made with the R package ‘tidyplots’, based on data show in Table S4. Gene presence is expressed as a percentage of ferns transcriptome containing an ortholog, out of 26 transcriptomes analysed. PTI: PAMP-triggered immunity, DAMP: Danger-associated molecular pattern, ETI: effector-triggered immunity, CNL: CC-NLR, TNL: TIR-NLR, SAR: systemic acquired resistance, AMP: antimicrobial peptide.

Proteins acting downstream of TIR-NLRs, such as EPs (EDS1, PAD4) and RPW8-NLRs (NRG1 and ADR1), are present in ferns but low copy number and divergent from their angiosperm counterparts (**Figure 5** and **7**) (Qin *et al*., 2024). This suggests that the EP/RPW8-NLR pathway regulating TIR-NLR immunity (Locci & Parker, 2024) is ancestral to streptophytes but was reduced in ferns.

In Arabidopsis, pathogen detection triggers the synthesis of salicylic acid (SA) that activates both local and systemic immune response, known as systemic acquired resistance (SAR) (Ding & Ding, 2020). SARD1 and CBP60g, that regulate the expression of genes for SA biosynthesis such as *ICS1* and *EDS5* (Wang *et al*., 2011), are strongly conserved in ferns (**Figure 7**). However, genes required for SA biosynthesis were not analysed by (Ali *et al*., 2024). NPR1, an SA receptor required for SAR in Arabidopsis (Dong, 2004), is mostly absent from ferns (**Figure 7**). In *Marchantia polymorpha*, NPR1 is conserved, can complement Arabidopsis *npr1* mutant and can bind SA, suggesting that this function is ancestral to land plants (Jeon *et al*., 2024). The absence of *NPR1* in many ferns questions the role of SA in SARD1/CBP60g-mediated immunity and SAR in ferns, and whether alternative mechanisms exist. Pipecolic acid is another molecule required for SAR, and its biosynthesis requires SARD4 (Ding *et al*., 2016). Unlike *NPR1*, *SARD4* is conserved in ferns, suggesting that pipecolic acid may play a predominant role in fern SAR.

Additionally, antimicrobial peptides (AMP), including chitinases and defensin, are widely present in ferns (**Figure 7**). It supports the idea that AMPs constitute a conserved defence strategy across eukaryotes (Zasloff, 2002; Zhu, 2007; Kulaeva *et al*., 2020).

Overall, many components of PTI and ETI appear to be ancestral to tracheophytes. However, some pathways, such as the EP-mediated regulation of TIR-NLR signalling, were reduced in ferns, suggesting the presence of yet-to-be-identified compensatory mechanisms.

### Ferns can be a reservoir of *R*-genes

The plant immune system relies on RLK/RLP cell-surface receptors for PTI or NLR cytoplasmic receptors for ETI (Jones & Dangl, 2006; Ngou *et al*., 2022b; Jones *et al*., 2024). Many disease resistance strategies in crops involve the introgression of such immune receptors (van Esse *et al*., 2019). Since we found that ferns can resist pathogens (**Figures 1** and **2**), we propose that they could serve as natural reservoirs of immune receptors, to be transferred to crops.

We found that PTI components are largely conserved across land plants (**Figures 4** and **7**). This is corroborated by functional studies showing conservation of central PTI regulators such as LysM receptors (Yotsui *et al*., 2023; Teyssier *et al*., 2024), SERK receptors (Yan *et al*., 2024), CPK28 (Dou *et al*., 2024) or RBOH (Chu *et al*., 2023). This conservation indicates that RLK/RLP could be prioritized as candidate determinants for resistance to pathogen in ferns.

In addition, NLRs have evolved differently across plant lineage (**Figure 5**). In ferns, we found three major classes of NLRs: CC-, TIR-and RPW8-NLRs (**Figure 5**). Fern CC-NLRs belong to the ‘Cbl-N’ cluster (**Figure 5A**), which is found in in bryophytes, lycophytes and ferns, but is lost in angiosperms, suggesting that ‘Cbl-N’ represent the ancestral state of CC-NLRs. Instead, angiosperms evolved ‘G10’ and ‘EDVID’ CC-NLRs. ‘G10’ and ‘EDVID’ expanded (**Figure 5**), to become overrepresented in asterids and monocots (Shao *et al*., 2016). Their success in asterids could be explained by the robustness of the ‘NRC’ networks, formed between many CC-NLRs ‘EDVID’ (Upson *et al*., 2018; Wu *et al*., 2018; Adachi & Kamoun, 2022; Goh *et al*., 2024). TIR-NLRs and RPW8-NLRs are found in all lineages (**Table S3**) (Qin *et al*., 2024). They have been shown to function in the same immune signalling pathway, along with the protein EDS1 (Castel *et al*., 2019; Feehan *et al*., 2020; Locci *et al*., 2023). This broad conservation suggests that it dates back to the earliest land plants. These data suggests that fern NLRs could be functional in angiosperm crops, despite millions of years of independent evolution. However, no fern NLR has yet been demonstrated to confer resistance to a pathogen.

Finally, we discovered a novel type of NLR, with a disordered N-terminal domain, which we called disN-NLR (**Figure 6**). disN-NLRs are mostly represented in ferns (**Figure 5B**) and are also present in lycophytes and bryophytes, but absent in angiosperms (**Table S3**, **Figure 6**).

In conclusion, we identified novel pathosystems in ferns, some of which being likely controlled by specific resistance mechanisms. We also found that fern genomes encode a remarkable diversity of putative immune receptors, including some absent in angiosperms. These findings suggest that ferns could serve as a reservoir of immune receptors for crop protection. Further characterisation of the newly discovered disN-NLRs could provide valuable insights into plant immune signalling and the evolution of disease resistance.

## ACKNOWLEDGMENTS

We are grateful to the Genotoul bioinformatics platform Toulouse Occitanie (Bioinfo Genotoul, https://doi.org/10.15454/1.5572369328961167E12) for providing computing resources. This study was supported by the Fédération de Recherche Agrobiosciences, Interactions et Biodiversité, the Laboratoires d’Excellence (LABEX) TULIP (ANR-10-LABX-41). This work was supported by the Agence Nationale de la Recherche (ANR LEVEL-UP ANR-21-CE20-0010-01) to C.J.

## COMPETING INTERESTS

None declared.

## AUTHOR CONTRIBUTIONS

B.C., C.J. and P.M.D. designed the research; B.C. and M.Ba. conducted the experiments; B.C., J.K. and M.Bo. analysed the data; B.C. and C.J. wrote the manuscript with input from all authors. All authors read and approved the final manuscript.

## DATA AVAILABILITY

All novel data and datasets presented in this study are available as main or supporting information.

## SUPPORTING INFORMATION

Figure S1: Phylogeny of ferns and microbes tested in this study

Figure S2: Wordclouds of fern RLK/RLP ‘others’ and disN-NLR predicted protein domains

Table S1: List of microbes, fern and control plant species genomes used in this study

Table S2: List of RLKs and RLPs from ferns and control species

Table S3: List of NLRs from ferns and control species

Table S4: Presence/absence of known immune genes (extracted from Ali et al, bioRxiv, 2024, table S9)

Data S1: Phylogeny of NB-ARCs (Newick)

Data S2: Phylogeny of NB-ARCs (colours)

Data S3: NLR sequences (FASTA)

Data S4: Alignment of disN-NLR n-terminal domain

Data S5: Phylogeny of disN-NLR n-terminal domain

Data S6: AlphaFold predicted structures of disN-NLRs

Method S1: Detailed Methods for NLR and RLK/RLP Mining

## SUPPLEMENTAL FIGURES

**Supplemental Figure 1:**
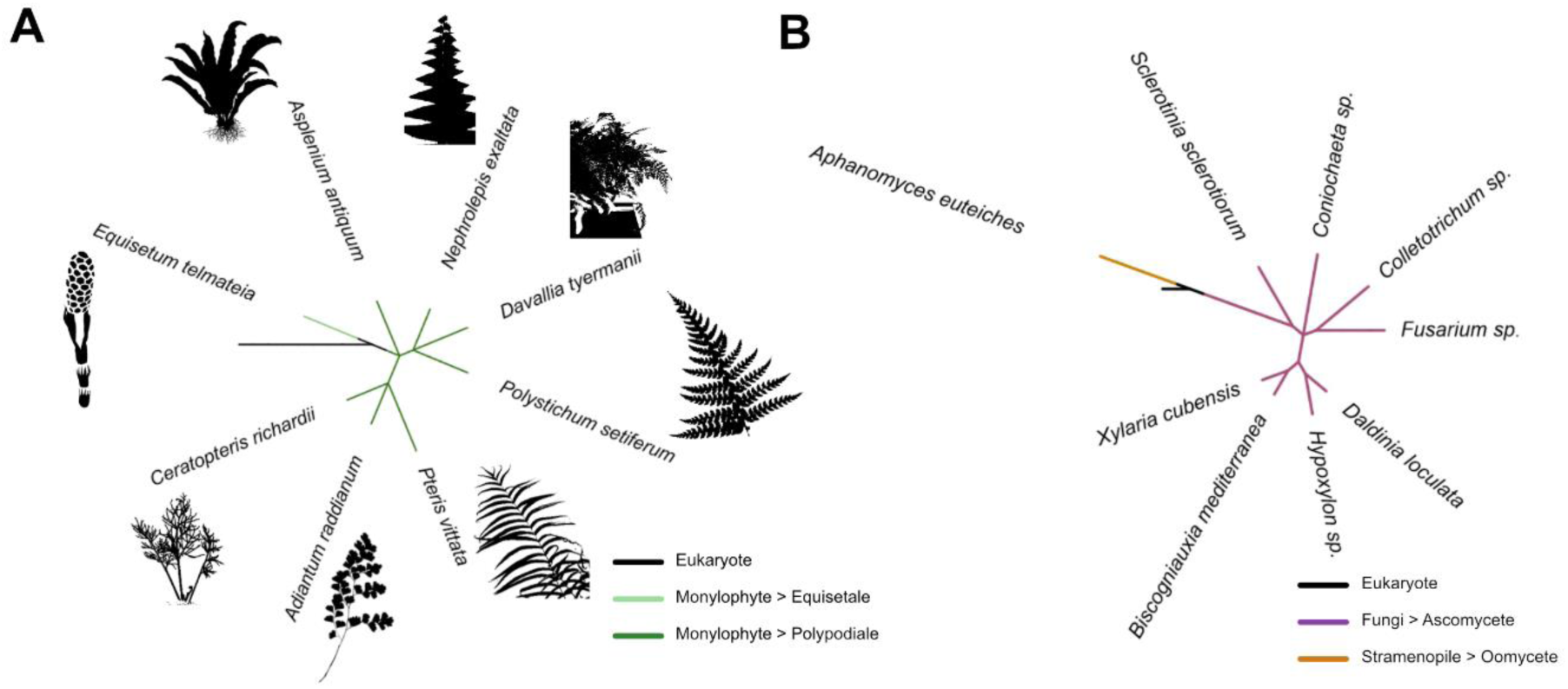
Phylogeny of ferns and microbes tested in this study. **A.** Phylogenetic tree of the fern species tested, generated using ‘phyloT’ (https://phylot.biobyte.de/) and visualised with iTOL (https://itol.embl.de/). The black branch represents eukaryotes, the light green corresponds to Equisetales and dark green to Polypodiales. Cartoons sources: *A. antiquum*, *A. raddianum*, *E. telmateia* (phylopics, https://www.phylopic.org), *C. richardii* (modified from https://c-fern.org/), *N. exalatata*, *P. setiferum*, *P. vittata* (this study). **B.** Phylogenetic tree of the microbial species tested, generated using ‘phyloT (https://phylot.biobyte.de/) and visualised using iTOL (https://itol.embl.de/). The black branch indicates eukaryotes, purple indicates opistokonts, orange represents stramenopiles.

**Supplemental Figure 2:**
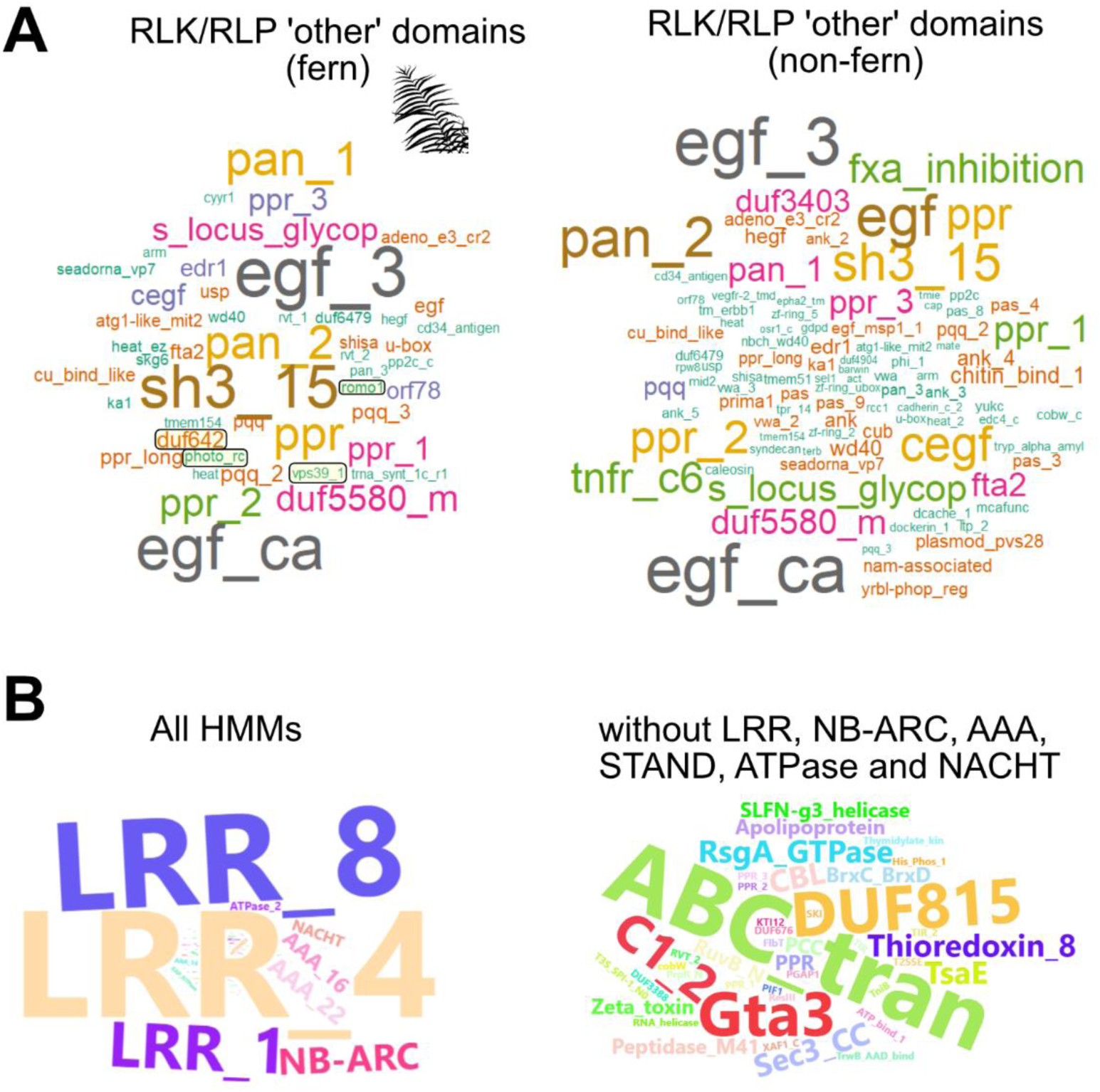
Wordclouds of fern RLK/RLP ‘others’ and disN-NLR predicted protein domains. **A.** Wordcloud representation of HMM domains detected in the ‘other’ category of RLKs/RLPs (mostly RLKs). Kinases and common signalling domains (such as EF-hand) were excluded for clarity. HMM domains unique to ferns are highlighted in black squares (romo1, vps39, mcm, photo_rc and duf642). Figure generated with the R package ‘wordcloud’. **B.** Wordcloud representation of HMM domains from disN-NLRs as presented in Figure 5. Left panel = all detected HMM domains, right panel = all HMM domains excluding LRR, NB-ARC, NACHT, ATPase, NTPase, STAND and AAA for imprived visibility. Figure generated with the R package ‘wordcloud2’.

## Notes

### Competing Interest Statement

The authors have declared no competing interest.

