## Supplementary material for "Exploring fern pathosystems and immune receptors to bridge gaps in plant immunity": Method_supp.docx

**Method S1: Detailed Methods for NLR and RLK/RLP Mining**

Immune receptors were predicted across available fern genomes and representative species from other land plant lineages (**Table S1**). Protein mining was performed using RGAugury v2.2 (Li *et al.*, 2016) and NLRtracker v0fe62b3 (Kourelis *et al.*, 2021) with default parameters.

For analyses on putative cell-surface immune receptors, proteins annotated as RLK or RLP by RGAugury were extracted (**Table S2**). The different RLK/RLPs were assigned to a family using keyword-based classification. Specifically, proteins were assigned to the LRR category if they contained ‘lrr’ (excluding proteins that also contained ‘lectin’), to the Lectin category if they contained ‘lectin’ (excluding those that also contained ‘lrr’), to the Malectin category if they contained ‘malectin’ (excluding those that also contained ‘lrr’), to the LRR-Malectin (LRR-Mal.) category if they contained both ‘lrr’ and ‘malectin,’ to the WAK category if they contained ‘wak,’ to the LysM category if they contained ‘lysm,’ to the GH category if they contained ‘glyco_hydro,’ to the Thaumatin category if they contained ‘thaumatin,’ to the SPARK category if they contained ‘spark,’ to the CRR category if they contained ‘stress-antifung,’ and to the RCC1 category if they contained ‘rcc1.’ RLK/RLPs that did not match any of these categories were assigned to the ‘other’ category (**Table S2**). To further investigate unclassified RLK/RLPs, domain composition was examined using HMM domain enrichment analysis, and wordcloud representations of domain composition were generated using the ‘wordcloud’ package in R (**Figure S2A**).

For analyses on putative intracellular receptors, 20,928 proteins selected by both NLRtracker and RGAugury were subjected to de novo functional domain annotation using HMMSCAN from the HMMER v3.2.1 package with an e-value threshold of 1^e-04^. Only proteins containing a predicted NB-ARC domain were retained, yielding 4,764 NB-ARC domains. These sequences were aligned using MUSCLE v5.1.0 with the ‘super5’ option enabled, and the alignment was trimmed using trimAl v1.4.1 (Capella-Gutiérrez *et al.*, 2009) to remove positions with >60% gaps. Phylogenetic reconstruction was performed using maximum likelihood with the IQ-TREE v2.2.2.6 program (Minh *et al.*, 2020), and the best-fitting evolutionary model was selected using ModelFinder (Kalyaanamoorthy *et al.*, 2017) according to the Bayesian Information Criteria. Branch support was estimated with 10,000 replicates of both sh-aLRT (Guindon *et al.*, 2010) and ultrafast bootstrap (Minh *et al.*, 2013). The resulting tree was visualized using iTOL v7 online platform.

To classify NLRs, a keyword-based approach was applied using domain annotations from RGAugury, NLRtracker, and HMMSCAN (**Table S3**). NLRs were categorized as follows: TIR-NLRs were identified by the presence of ‘TIR,’ ‘TN,’ or ‘TNL’ but not ‘TPR.’ CC-NLRs were identified by the presence of ‘CC,’ ‘Rx_N,’ ‘CNL,’ or ‘CN’ but not ‘arm.’ RPW8-NLRs were identified by the presence of ‘RPW8’ or ‘CCR.’ Hyd-NLRs were identified by the presence of ‘αβhydrolase,’ ‘a/b,’ or ‘DUF676.’ Kin-NLRs were identified by the presence of ‘Pkin’ or ‘PK_Tyr_Ser-Thr.’ TIR-NB-TPRs were identified by the presence of both TIR/TNL and TPR domains. CC-NB-ARMs were identified by the presence of both CC/CNL and arm domains. Proteins that did not fit into any of these categories were labeled as ‘Undetermined’ (**Table S3**).

Branches in the phylogenetic tree that were well-supported (bootstrap >70) were manually color-coded using iTOL: yellow for TIR-NLRs, red for Hyd-NLRs, purple for Kin-NLRs, blue for CC-NLRs, green for RPW8-NLRs, mustard for TIR-NB-TPRs, and dark blue for CC-NB-ARCs (**Figure 5A**, **Data S1** and **S2**). Some NLRs did not fit into known classifications. Five CC-NLRs from *Camellia sinensis* clustered with TIR-NLR ‘branch III’ (blue). Two NLRs grouped with Hyd-, Kin-, and TIR-NLRs (branches I and II, black). Eight NLRs from mosses clustered with Kin-NLRs but lacked a kinase domain. Fourteen angiosperm NLRs formed a branch with TIR-NB-TPRs. Five TIR-NLRs clustered with CC-NLRs (yellow).

The final dataset consisted of 3,775 NB-ARC domains from 3,637 proteins. The tree is available n newick format (**Data S1**). The colour annotation are stored in TXT format (**Data S2**). Full-length NB-ARC protein sequences are provided in FASTA format (**Data S3**).
