## Supplementary figures and images for "Exploring fern pathosystems and immune receptors to bridge gaps in plant immunity"

### FigS1.png

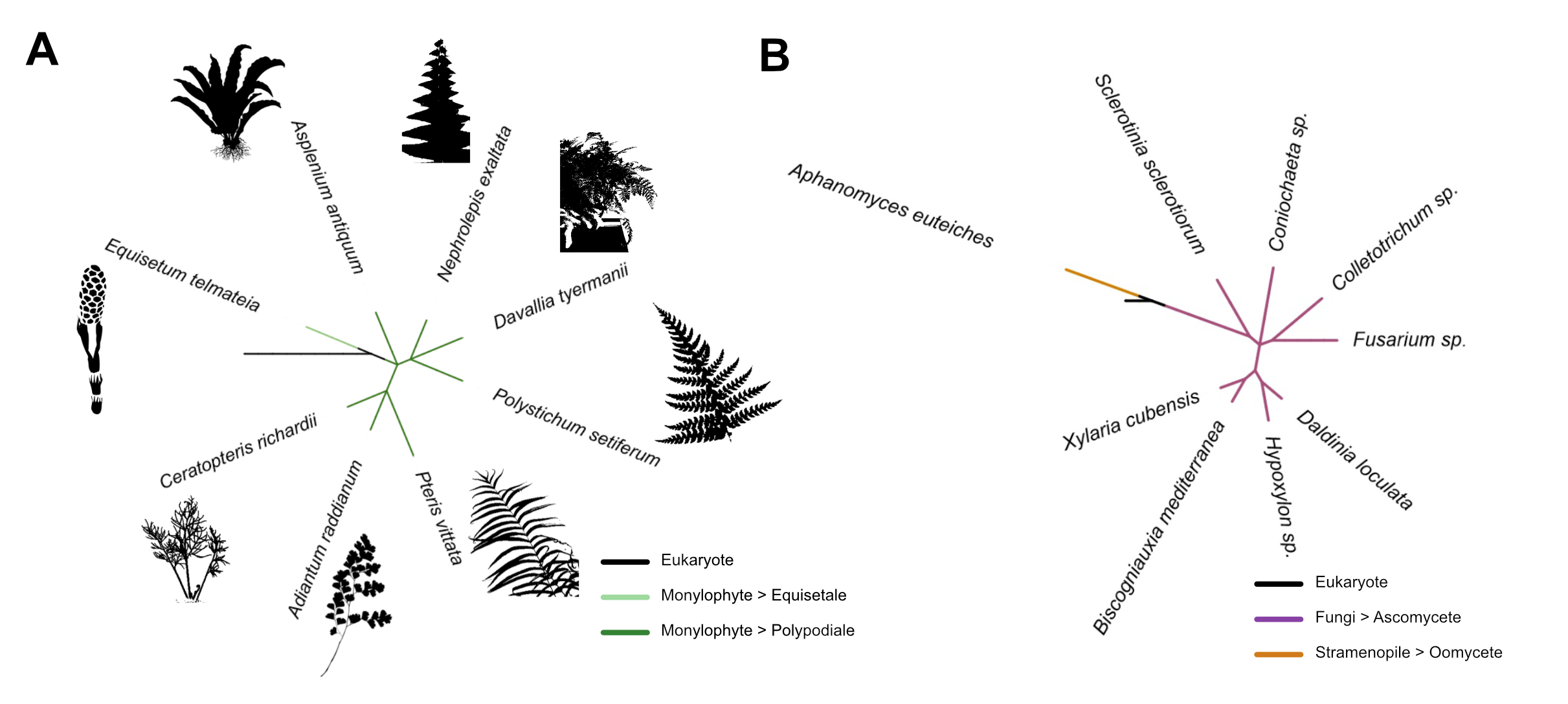

### FigS2.png

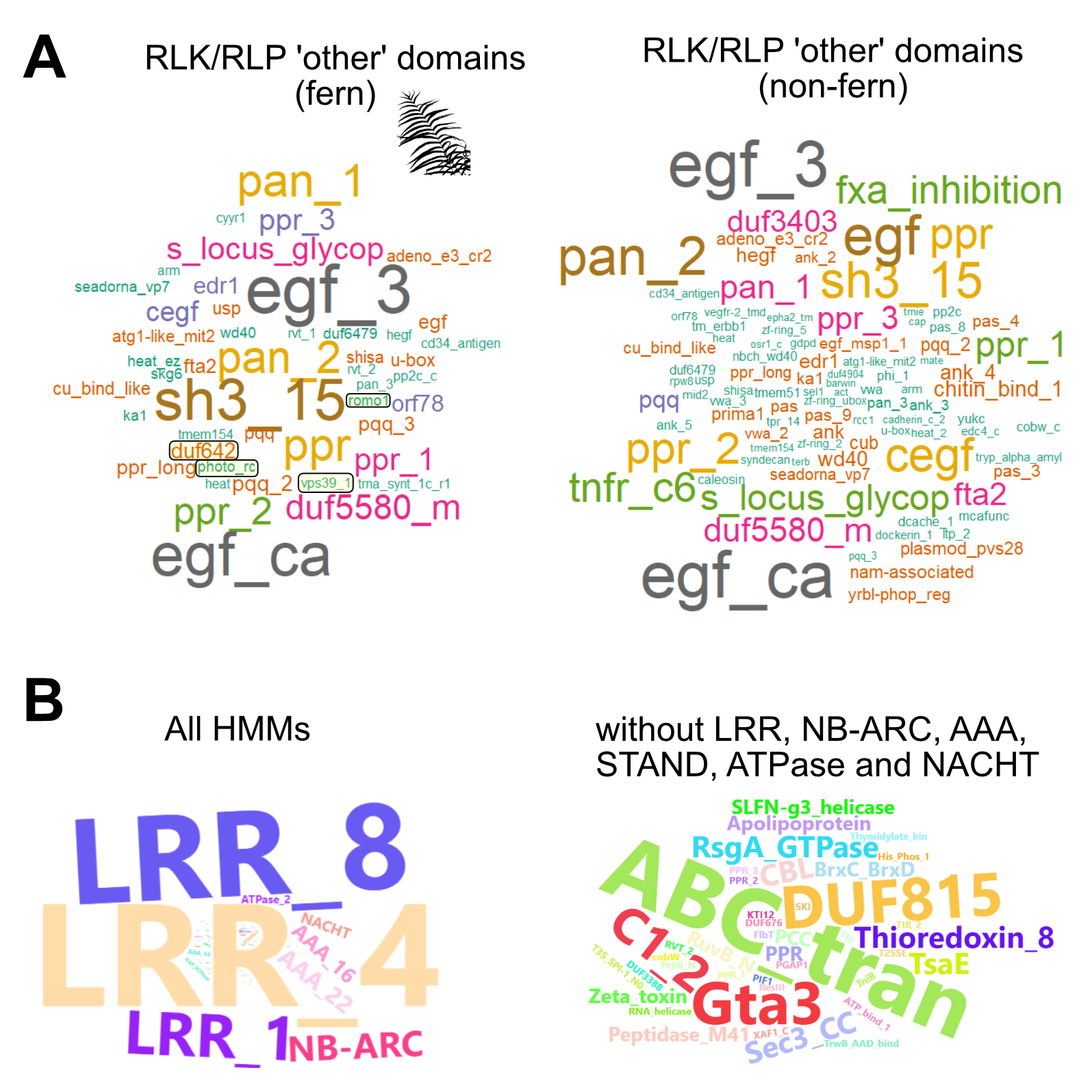
